# The circadian regulator PER1 inhibits osteoclastogenesis by activating inflammatory genes

**DOI:** 10.1101/2025.09.18.677145

**Authors:** Nobuko Katoku-Kikyo, Elizabeth K. Vu, Samuel Mitchell, Ismael Y. Karkache, Elizabeth W. Bradley, Nobuaki Kikyo

## Abstract

Disruption of circadian rhythms predisposes shift workers to many chronic conditions, including osteoporosis. However, the effects of disrupted circadian rhythms on bone remodeling remain largely unknown. Here, we show that one of the core circadian regulators PER1 inhibits osteoclastogenesis by upregulating genes involved in inflammation. The conditional knockout of *Per1* in osteoclasts and related cells resulted in decreased bone mass in the femurs of mice, along with increased osteoclasts and decreased osteoblasts. Osteoclastogenesis was also promoted by *Per1* depletion in vitro with 16 downregulated inflammatory genes. Seven of these genes were known to promote or inhibit osteoclastogenesis depending on the stage of osteoclastogenesis and the presence or absence of infection. Knockdown of *Nlrp3*, *Tlr8*, or *Tlr9* in the group of genes promoted osteoclastogenesis, mirroring the effects of *Per1* knockout and offering a mechanistic explanation for the *Per1*-mediated inhibition of osteoclastogenesis. These results were not observed following the knockout of a paralog *Per2*. *Per1* knockout mice maintain general circadian rhythms, unlike arrhythmic *Per1;Per2* double knockout mice. This gives credence to *Per1* as a selective target for therapeutic interventions without disrupting the circadian rhythms. This study uncovered a molecular link between a circadian regulator and osteoclastogenesis in the broader context of inflammatory reactions. Our findings may be mechanistically relevant to inflammatory bone diseases influenced by circadian rhythms, such as rheumatoid arthritis and osteoarthritis, as well as other bone diseases predisposed by chronic circadian disruption.

**Lay summary:** Disruption of circadian rhythms is a risk factor for many chronic diseases, including osteoporosis, among shift workers; however, underlying mechanisms remain largely unknown. In this study, the depletion of *Per1*, a core circadian regulator, resulted in an increase in bone-resorbing osteoclasts and a decrease in bone mass in mice. These changes were accompanied by a decrease in the expression of inflammatory genes that promoted the formation of osteoclasts upon depletion. This study revealed a link between circadian rhythms and bone loss, with inflammatory genes serving as mediators, which could provide a basis for future therapeutic interventions.

## Introduction

Disruption of circadian rhythms due to frequent travel, shift work, or excessive exposure to artificial light increases the risk of osteoporosis,^1,2^ a condition characterized by an imbalance in bone formation by osteoblasts and bone resorption by osteoclasts, both of which are under circadian control^3^. Mammalian circadian rhythms are regulated by the central clock located in the suprachiasmatic nucleus of the hypothalamus and peripheral clocks ubiquitously expressed throughout the body.^4,5^ The central clock is entrained by light signals transmitted from the retina and synchronizes peripheral clocks via the autonomic nervous system and various hormones. Peripheral clocks are also entrained by other stimuli, such as food intake and sleep-awake signals. All these clocks are maintained by two feedback loops centered on the CLOCK/BMAL1 transcription factor complex. This heterodimer binds to the E-box in hundreds of target genes and activates their transcription. The target genes include *Cry* (*Cry1* and *Cry2*) and *Per* (*Per1*, *Per2*, and *Per3*), which form the CRY-PER heterodimer and inhibit CLOCK/BMAL1 through a direct interaction. Subsequently, both CRY and PER are degraded via the proteasomal pathway, enabling CLOCK/BMAL1 to resume transcription of the target genes and completing the first feedback loop with a 24-hr period. In the second feedback loop, CLOCK/BMAL1 activates the transcription of the *Ror* (retinoic acid receptor-related orphan receptor) and *Nr1d* (Rev-erbs or reverse orientation c-erbA) genes, whose proteins then bind to the *Bmal1* promoter and activate (ROR) or inhibit (NR1D) transcription.

Circadian regulation of bone remodeling has been primarily demonstrated through three lines of evidence.^3,6^ First, serum and urine levels of bone turnover markers representing bone resorption (such as crosslinked C-terminal telopeptide of type I collagen [CTX]) and bone formation (such as procollagen type 1 N-terminal propeptide [P1NP]) exhibit circadian rhythms. Second, RNA-seq and quantitative reverse transcription and PCR (qRT-PCR) demonstrate circadian expression of bone marker genes, such as *Runx2*, *Nfatc1*, *Ctsk*, *Tnfrsf11a* (*Rank)*, and *Tnfrsf11b* (*Opg*) in the mouse calvaria, tibia, and femur. Third, bone cell-specific knockout (KO) of circadian regulators disrupts bone mass. For example, osteoblast-specific KO of *Bmal1* decreases bone mass by modulating the RANKL signaling pathway.^7^ In contrast, osteoclast-specific KO of *Bmal1* increases bone mass,^8^ although another study was unable to replicate this finding.^9^ However, since the depletion of *Bmal1* completely abolishes circadian rhythms, these phenotypes could include general consequences of the lack of circadian rhythms, in addition to the intended *Bmal1-*specific effects.^10^ The roles of non-*Bmal1* circadian regulators in osteoclastogenesis remain poorly understood.

We depleted *Per1* or *Per2* in the osteoclasts of mice in the current study for two reasons. First, single-gene germline KO mice for either gene can maintain circadian rhythms with a period of approximately 22 hr, unlike arrhythmic double KO mice,^11^ addressing the problem mentioned above. Although germline KO of *Per1* or *Per2* does not result in an overt skeletal phenotype,^12^ this does not exclude hidden roles of the genes in osteoclastogenesis. Second, *Per1* KO promotes the production of pro-inflammatory cytokines in macrophages under inflammatory conditions as we reported before^13^, suggesting that *Per1* KO might affect osteoclasts as well because of the shared lineage between macrophages and osteoclasts.

In this study, we used *Cx3cr1* promoter-driven *Cre* mice to conditionally KO (cKO) *Per1* or *Per2* in the monocyte/macrophage lineages. We first compared the bone phenotypes of *Per1* cKO and *Per2* cKO mice using micro-computed tomography (micro-CT) and histology. We then used in vitro osteoclastogenesis to verify the cell-autonomous effects of *Per1* cKO. Subsequently, transcriptomics, gene knockdown (KD), and circadian synchronization led us to identify a group of genes that could explain the phenotypes of *Per1* cKO. Collectively, these results revealed inflammatory gene activation as a major link between *Per1* and osteoclasts.

## Materials and Methods

### Generation of *Per1* and *Per2* cKO mice

*Per1* cKO mice were created by crossing the *Cx3cr1* promoter-driven *Cre* mice (B6J.B6N(Cg)-*Cx3cr1^tm1.1(cre)Jung^*/J, Jackson Lab, 025524) with the inducible *Per1* KO mice *Per1^fl/fl^*.^14^ The resulting *Cx3cr1-Cre^tg/wt^;Per1^fl/fl^*and *Cx3cr1-Cre^wt/wt^;Per1^fl/fl^* mice were used as *Per1* cKO and littermate control mice (*Per1* Cont), respectively. Likewise, *Cx3cr1*-*Cre* mice and *Per2^fl/fl^*mice (Medical Research Council, UK, Per2^tm1c(EUCOMM)Hmgu^) were crossed to establish *Cx3cr1-Cre^tg/wt^;Per2^fl/fl^* and *Cx3cr1-Cre^wt/wt^;Per2^fl/fl^* mice as *Per2* cKO and littermate control mice (*Per2* Cont), respectively. Genotyping primers and the sizes of the PCR products are listed in **Supplementary Table 1** based on the protocols provided by the sources of the mice. The mice were housed under a 12 hr light-12 hr dark cycle in an accredited facility with water and food provided *ad libitum*. All mouse protocols were approved by the Institutional Animal Care and Usage Committee of the University of Minnesota (2410-42491A). All mouse experiments comply with the standards set by the Guide for the Care and Use of Laboratory Animals published by the US National Institute of Health.

### Micro-CT scan analysis of the bones

Micro-CT scanning of the femur was performed as previously described.^15^ Specifically, the femurs prepared from 12-week-old mice were fixed in 10% formalin in phosphate-buffered saline (PBS) for 24 hr and stored in 70% ethanol at 4°C. Micro-CT was performed with an XT H 225 CT scanning system (Nikon Metrology). The scan settings were as follows: 120 kV, 61 μA, 720 projections at two frames per projection with an integration time of 708 ms, an isometric voxel size of 7.11 μm with a 1 mm aluminum filter, and 17 min per scan. Bone morphometric analysis was done with SkyScan CT Analyzer (CTAn, Brucker micro-CT) and the results were described with standard symbols.^16^ Each scan volume was reconstructed with CT Pro 3D program (Nikon Metrology), converted to bitmap data with VGStudio MAX 3.2 (Volume Graphics), and reoriented with DataViewer (SkyScan, Buker micro-CT) for qualitative analysis. 3D analysis of trabecular bones was done at the distal metaphysis 0.7 mm proximal to the growth plate, extending 1.5 mm toward the bone diaphysis. 3D analysis of the cortical bones was done in a 0.5 mm section in the mid-diaphysis 4 mm from the growth plate. Ten femurs were collected in each group and none of them were excluded in the analysis. Bone samples were blinded but not randomized during analysis. The G*Power program (https://download.cnet.com/g-power/3000-2054_4-10647044.html#google_vignette) was used in the power analyses of this and the histomorphometry below with the significance level of 0.05 and the power value of 0.8.

### Histomorphometry of bone sections

The tibiae prepared as described above were decalcified in 15% EDTA for 14 days, embedded in paraffin, and sectioned at 7 μm in thickness. The TRAP staining kit (Sigma Aldrich, 387A-1KT) and Masson’s trichrome staining kit (Polyscience, 25088) were used to detect osteoclasts and osteoblasts, respectively. The numbers of osteoclasts and osteoblasts were counted and the ratios between these cells and bone perimeter were calculated with ImageJ. Seven tibiae were randomly selected and all of them were analyzed without exclusion. Sample blinding was not applied.

### Osteoclast differentiation in vitro

Osteoclasts were prepared from bone marrow monocytes and macrophages (BMMs) as previously described with some modifications.^17^ On day 1, bone marrow cells were flushed from the femur and tibia of 8- to 12-week-old mice with PBS and centrifuged at 190 x *g* for 5 min. The precipitated cells were resuspended with 1X Red Blood Cell Lysis Buffer (eBioscience, 00-4333-57) and incubated at 25°C for 5 min. After centrifugation under the same conditions, the precipitated cells were resuspended with Monocyte Medium (25 ng/ml M-CSF [Shenandoah, 100-03], 10% heat-treated fetal bovine serum [FBS], and phenol red-free αMEM [Thermo Fisher, 41061-029]). Cells were incubated with 5% CO2 at 37°C overnight. On day 2, non-adherent cells were harvested and cell clusters were removed with a 70 μm cell strainer (Falcon, 352350). The cells were centrifuged, resuspended in Osteoclast Medium (Monocyte Medium with 100 ng/ml RANKL [Cell Signaling Technology, 68495]), and seeded at 8x10^5^ cells/400 μl/well in a 48-well plate. On day 4, 200 μl of the medium was added. On day 5, the medium was replaced with 400 μl of fresh medium. On day 6, the cells were fixed with 4% paraformaldehyde in PBS for TRAP staining. The total number of osteoclasts, defined as TRAP-positive cells with more than two nuclei, in a well were manually counted. The sizes of osteoclasts were measured with ImageJ.

### Enzyme-linked immunosorbent assay (ELISA) of IL-1β and lipopolysaccharide (LPS)

Culture supernatant of osteoclasts on day 6 were applied to ELISA of IL-1β (Lumit IL-1β immunoassay, Promega, W7010) and LPS (ToxiSensor chromogenic LAL endotoxin assay kit, GenScript, L00350Y) following each instruction.

### Synchronization of circadian rhythms

On day 5 of the differentiation of osteoclasts, the cells were treated with Synchronization Medium (50% heat-inactivated horse serum in Osteoclast Medium instead of FBS) for 1 hr at 37°C with 5% CO2, washed with PBS at 37°C twice, and cultured with fresh Osteoclast Medium.^18^ The time of completion of these procedures was defined as 0 hr post-synchronization. The cells were cultured at 37°C with 5% CO2 and harvested for qRT-PCR every 4 hr for 48 hr starting from 24 hr post-synchronization to wait for the recovery from the synchronization as previously recommended.^19^ Circadian rhythmicity, amplitude, and period of the qRT-PCR values was evaluated with harmonic regression using Cosinor.Online (https://cosinor.online/app/cosinor.php).^20^

### Bone resorption assay

The bone resorption assay followed a published protocol with some modifications.^17^ On day 2 of the osteoclast differentiation, the cells were seeded on top of a bone slice (Immunodiagnostic Systems, DT-1BON1000-96) at 3.2x10^5^ cells/well in a 48-well plate in Osteoclast Medium to induce osteoclastogenesis as described above. The medium was replaced every 3 days after day 6. On day 14, cells were lysed with 10% bleach and the bone slices were washed with deionized water three times. The bone slices were incubated in 200 μl deionized water for 1 hr at 25°C twice and stained with Toluidine blue for 30 sec. The bone slices were washed with deionized water three times and dried. The resorption area was analyzed with ImageJ.

### qRT-PCR

qRT-PCR was performed as previously described.^13^ RNA was extracted from cells using a Quick RNA Microprep kit (Zymo Research, R1051) and purity was assessed using a microvolume spectrophotometer DS-11 FX+ (Denovix). cDNA was synthesized with ProtoScript II Reverse Transcriptase (New England Biolabs, M0368L). qPCR was performed with the primers listed in **Supplementary Table 2** and qPCRBIO SyGreen Blue Mix Lo-ROX (Genesee Scientific, 17-505B) on a Mastercycler realplex^2^ thermocycler (Eppendorf). PCR conditions were as follows: initial denaturation at 95°C for 2 min, 40 cycles of 95°C for 5 sec - 60°C for 30 sec - 72°C for 30 sec, and a melting curve step to check the specificity of the amplification. mRNA expression levels were analyzed by normalizing expression values to glyceraldehyde 3-phosphate dehydrogenase (*Gapdh*) expression. Mean ± SEM of biological triplicates with technical triplicates each was calculated.

### Gene KD with siRNA

BMMs were transfected with 10 nM siRNA (**Supplementary Table 3**) with 1 ul DharmaFECT 1 Transfection Reagent (Dharmacon, T-200-01) for 5 hr on day 4 and 5 during osteoclast differentiation described above. The cells were harvested on day 6 for qRT-PCR and TRAP staining.

### Transfection of circadian genes into RAW264.7 cells

RAW264.7 cells (ATCC, TIB-71) were transfected with empty vector or plasmids encoding *Clock*, *Bmal1*, *Per1*, and *Per2* as follows.^21^ Cells were seeded at 8x10^4^ cells/well in a 48-well plate on day 1 in 10% FBS in DMEM. On day 2, 1.6 μg of the plasmid was transfected with 2 μl Lipofectamine LTX and 1.5 μl PLUS Reagent (Invitrogen, 15338030) following the instructions. The cells were incubated at 37°C with 5% CO2 and harvested 48 hr later for qRT-PCR.

### RNA-seq

RNA-seq and data analysis were also performed as previously described.^22^ Total RNA was prepared from day 6 osteoclasts and concentration and RNA integrity number were quantified with an Agilent BioAnalyzer 2100. mRNA was purified with poly-T oligo-attached magnetic beads and cDNA was synthesized using random hexamer primers. Non-directional libraries were prepared and completed by end repair, A-tailing, adapter ligation, size selection, amplification, and purification. The quality and quantity of the libraries were checked via real-time PCR and a Bioanalyzer. The libraries were sequenced on an Illumina platform and paired-end reads were generated.

Raw reads of the fastq format were processed through in-house perl scripts. Paired-end clean reads were aligned to the reference genome Mus musculus GRCm38 (ftp://ftp.ensembl.org/pub/release-94/fasta/mus_musculus/dna/Mus_musculus.GRCm38.dna.primary_assembly.fa.gz and ftp://ftp.ensembl.org/pub/release-94/gtf/mus_musculus/Mus_musculus.GRCm38.94.gtf.gz) using Hisat2 v2.0.5 (https://daehwankimlab.github.io/hisat2/). featureCounts v1.5.0-p3 (http://subread.sourceforge.net/) was used to count read numbers mapped to each gene. Differential expression analysis was performed using the DESeq2 R package (1.20.0) (https://www.r-project.org/). Genes with log2 fold change > 0.58 or < –0.58 (> 1.5-fold) and an adjusted p-value < 0.05 were assigned as differentially expressed. Enrichment of specific gene pathways in differentially expressed genes was identified by applying the clusterProfiler R package (https://www.r-project.org/) to the databases of Gene Ontology (http://www.geneontology.org). Adjusted p-value < 0.05 was considered as significant enrichment.

### Statistical Analysis

Unpaired two-tailed t-tests and two-way ANOVA with Tukey’s method of multiple comparisons were used as stated in the figure legends. Mean ± SEM was obtained from biological replicates of the numbers indicated in each figure. Box plots show median and interquartile range (25th–75th percentile). GraphPad Prism 10 (GraphPad Software) was used in statistical analysis.

## Results

### *Per1* cKO, but not *Per2* cKO, decreased bone mass and increased osteoclasts in male mice

To investigate the roles of *Per1* and *Per2* in osteoclasts, we prepared *Cx3cr1* promoter-driven *Per1* KO mice (*Per1* cKO) alongside littermate controls (*Per1* Cont), as well as *Per2* cKO mice with controls (*Per2* Cont). qRT-PCR verified significant depletion of *Per1* (18.3 ± 2.5% remaining) and *Per2* (20.7 ± 3.5% remaining) mRNA in osteoclasts prepared from the femurs of 12-week-old male mice (**Figure 1A**). Micro-CT analysis of the femoral midshaft revealed a significant reduction in cortical bone volume and cortical thickness in male *Per1* cKO mice compared to those in *Per1* Cont (**Figure 1B-D**). Likewise, diminished distal femoral trabecular bone volume and numbers were observed in *Per1* cKO mice (**Figure 1B**, **E**, and **F**). In contrast, *Per2* cKO mice did not show any of these phenotypes (**Figure 1B-F**). Trabecular thickness was not affected in *Per1* cKO or *Per2* cKO mice (**Figure 1G**). In this study, we focused on male mice because none of the bone mass parameters were decreased in female *Per1* cKO and *Per2* cKO mice compared to corresponding control mice (**Supplementary Figure 1A**-**E**).

**Figure 1.**
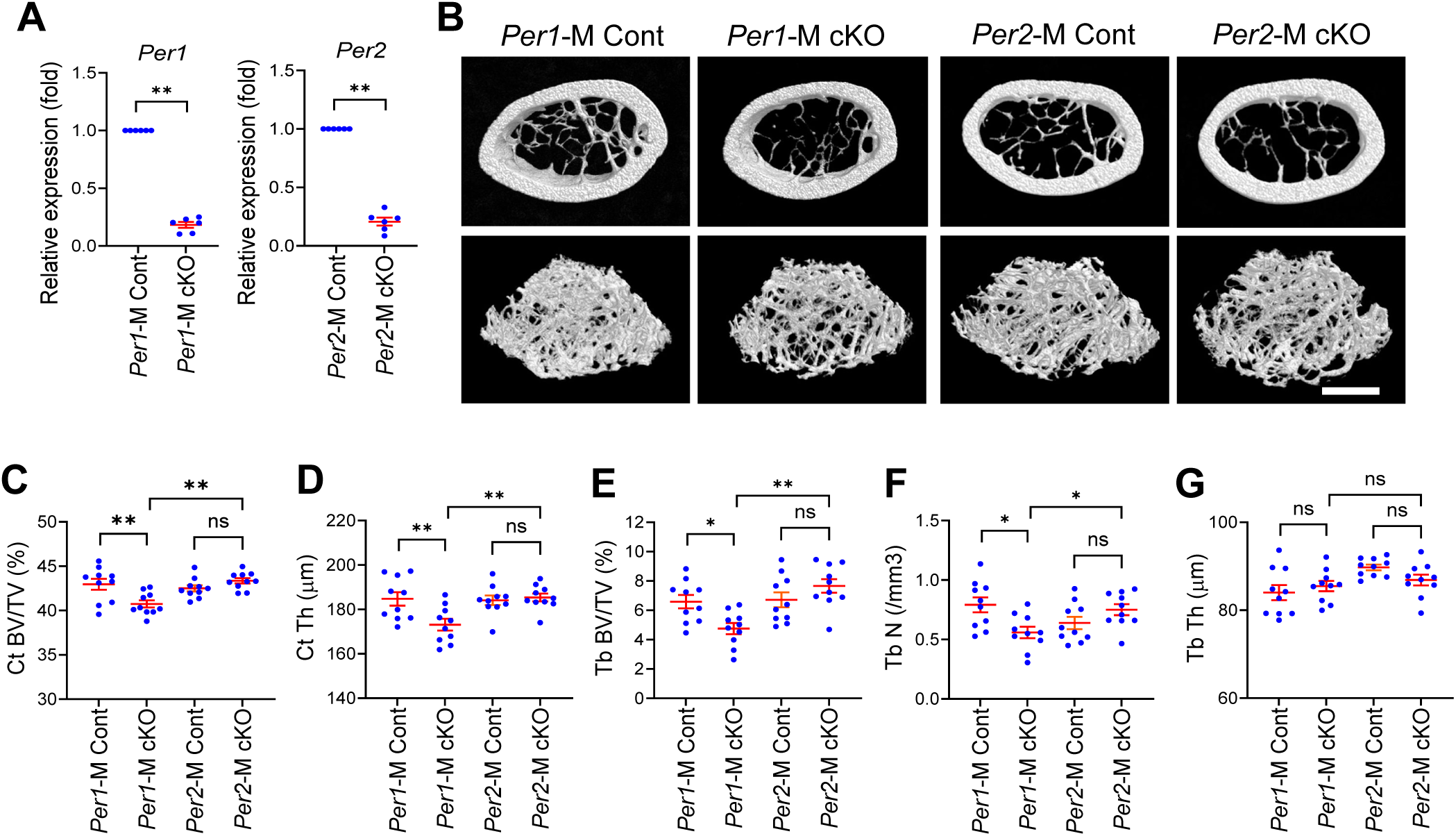
*Per1* cKO, but not *Per2* cKO, decreased bone mass in male femurs. **A.** Relative expression levels of *Per1* and *Per2* in each cKO male osteoclasts in comparison to Cont osteoclasts on day 6 quantified with qRT-PCR. n = 6. **B.** Reconstructed 3D micro-CT images of femoral midshafts (top) and distal femurs (bottom) prepared from 12-week-old male mice of the indicated genotypes. Bar, 1 mm. **C–G**. Quantification of the cortical bone volume/total volume ratio (**C**), cortical thickness (**D**), trabecular bone volume/total volume ratio (**E**), trabecular number (**F**), and trabecular thickness (**G**) comparing 12-week-old male mice. n = 10. Mean ± SEM is shown. ** p < 0.01, * p < 0.05, and ns for not significant with two-way ANOVA with Tukey’s method of multiple comparisons.

In bone histomorphometry of the proximal tibia, *Per1* cKO increased osteoclasts (**Figure 2A** and **B**) and reduced osteoblasts per bone perimeter (**Figure 2C** and **D**). The decrease in osteoblasts was likely an indirect effect because they do not express *Cx3cr1*.^23^ In contrast, the numbers of osteoclasts and osteoblasts were not changed following *Per2* cKO (**Figure 2B** and **D**, and **Supplementary Figure 2A** and **B**). The changes in the numbers of osteoclasts and osteoblasts in *Per1* cKO mice were consistent with the decreased bone mass in these mice.

**Figure 2.**
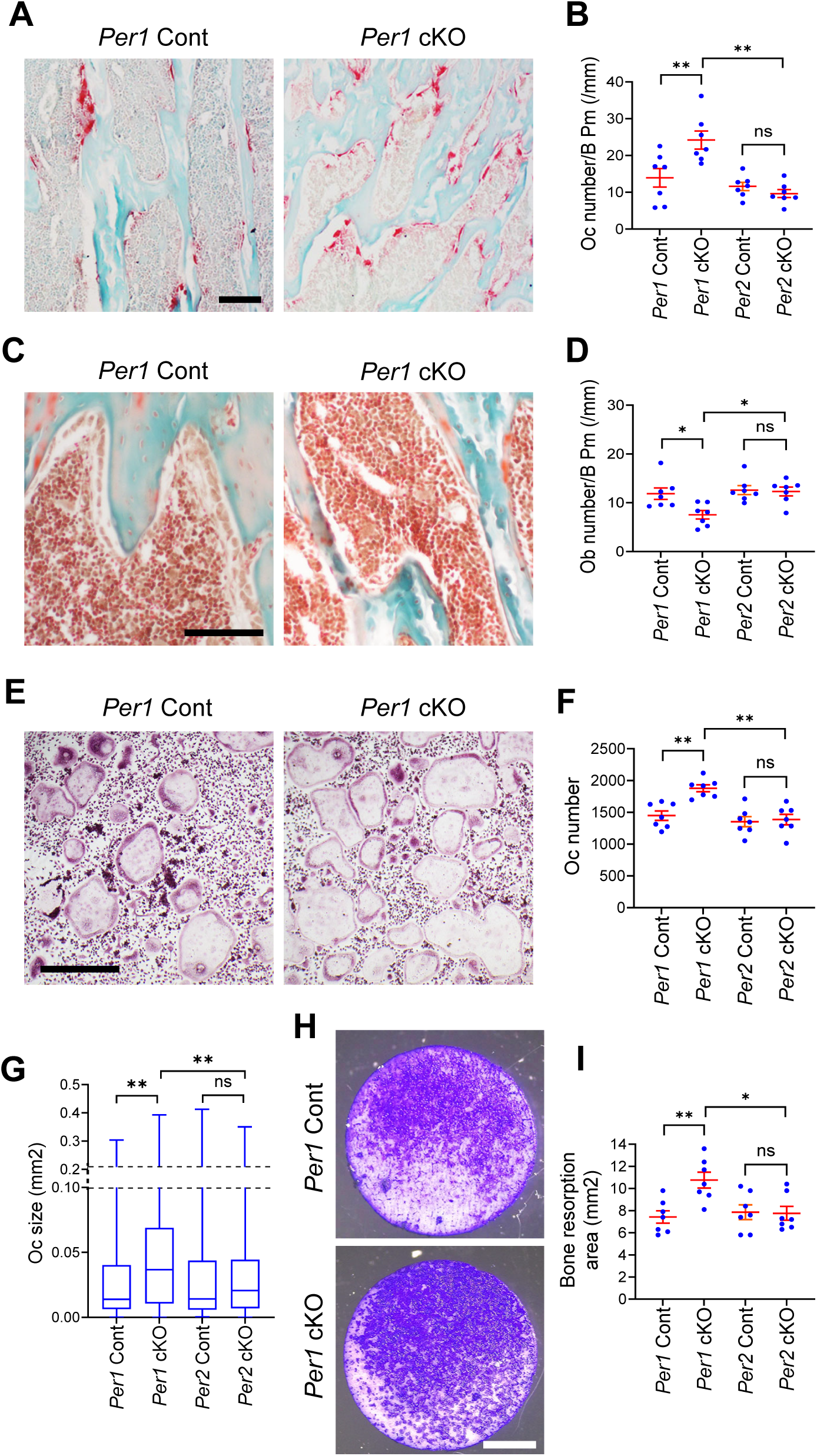
*Per1* cKO male mice showed increased osteoclastogenesis and decreased osteoblastogenesis. **A.** TRAP staining of the proximal tibial sections comparing *Per1* cKO and Cont mice. Bar, 100 μm. **B.** Osteoclast numbers per bone perimeter in the proximal tibiae of the indicated genotypes. n = 7. **C.** Masson’s trichrome staining of the proximal tibial sections. Bar, 100 μm. **D.** Osteoblast numbers per bone perimeter in proximal tibiae of the indicated genotypes. n = 7. **E.** TRAP staining of osteoclasts in vitro on day 6. Bar, 500 μm. **F.** The numbers of osteoclasts per well in a 48-well plate on day 6. n = 7. **G.** Size distributions of osteoclasts on day 6. n = 4,200 osteoclasts (600 osteoclasts/well x 7 wells of biologically independent experiments). **H.** Bone resorption assay stained with Toluidine blue. Bar, 2 mm. **I.** Areas of resorbed bones stained with Toluidine blue. n = 7. Histological sections and osteoclasts were prepared from 12-week-old male mice. Mean ± SEM is shown. ** p < 0.01, * p < 0.05, and ns for not significant with two-way ANOVA with Tukey’s method of multiple comparisons.

### *Per1* cKO promoted osteoclastogenesis in a cell-autonomous manner

We investigated whether *Per1* depletion increased osteoclastogenesis in a cell-autonomous manner, as the *Cx3cr1* promoter was also active in monocytes and macrophages. We harvested BMMs from the femurs of 12-week-old *Per1* cKO and Cont mice and induced osteoclastogenesis with RANKL and M-CSF. Greater numbers of large-sized osteoclasts were obtained from *Per1* cKO cells compared to those from Cont cells (**Figure 2E-G**). Moreover, enhanced pit formation was observed on bovine bone slices caused by *Per1* cKO osteoclasts (**Figure 2H** and **I**). These phenotypes were not detected with *Per2* cKO osteoclasts (**Figure 2F**, **G**, and **I**, and **Supplementary Figure 2C** and **D**). These in vitro results align with the observed decrease in bone mass and increase in osteoclasts in vivo.

### Inflammatory genes were downregulated in *Per 1* cKO osteoclasts

We applied RNA-seq to compare transcriptomes of *Per1* KO and Cont osteoclasts; 25 genes were upregulated and 96 were downregulated in *Per1* cKO osteoclasts when cutoff values of > 1.5-fold difference (log2FC > 0.58 or < −0.58) and an adjusted p-value of < 0.05 (−log10(padj) > 1.3) were applied (**Figure 3A** and **B**, **Supplementary Figure 3** and **4A**, and **Supplementary Table 4**). *Per1* and *Per2* were not downregulated in each cKO cell because residual exons generated mRNA. Gene ontology (GO) analysis of the upregulated genes did not yield any enriched pathways; however, we observed enrichment of downregulated genes involved in innate and adaptive immunity, as well as inflammatory response (**Figure 3C**), which included 16 genes shown in **Figure 3D** (red bars for *Per1* cKO). RNA-seq analysis of *Per2* cKO osteoclasts identified 2 upregulated and 30 downregulated genes compared to *Per2* Cont cells (**Supplementary Figure 4B** and **C**, **5A** and **B**, and **Supplementary Table 4**). However, there were no enriched pathways in GO analysis of these genes. In addition, the 16 genes mentioned above were not downregulated in *Per2* cKO osteoclasts (**Figure 3D**, blue bars), suggesting specific relevance of these genes to the increased osteoclastogenesis associated with *Per1* cKO.

**Figure 3.**
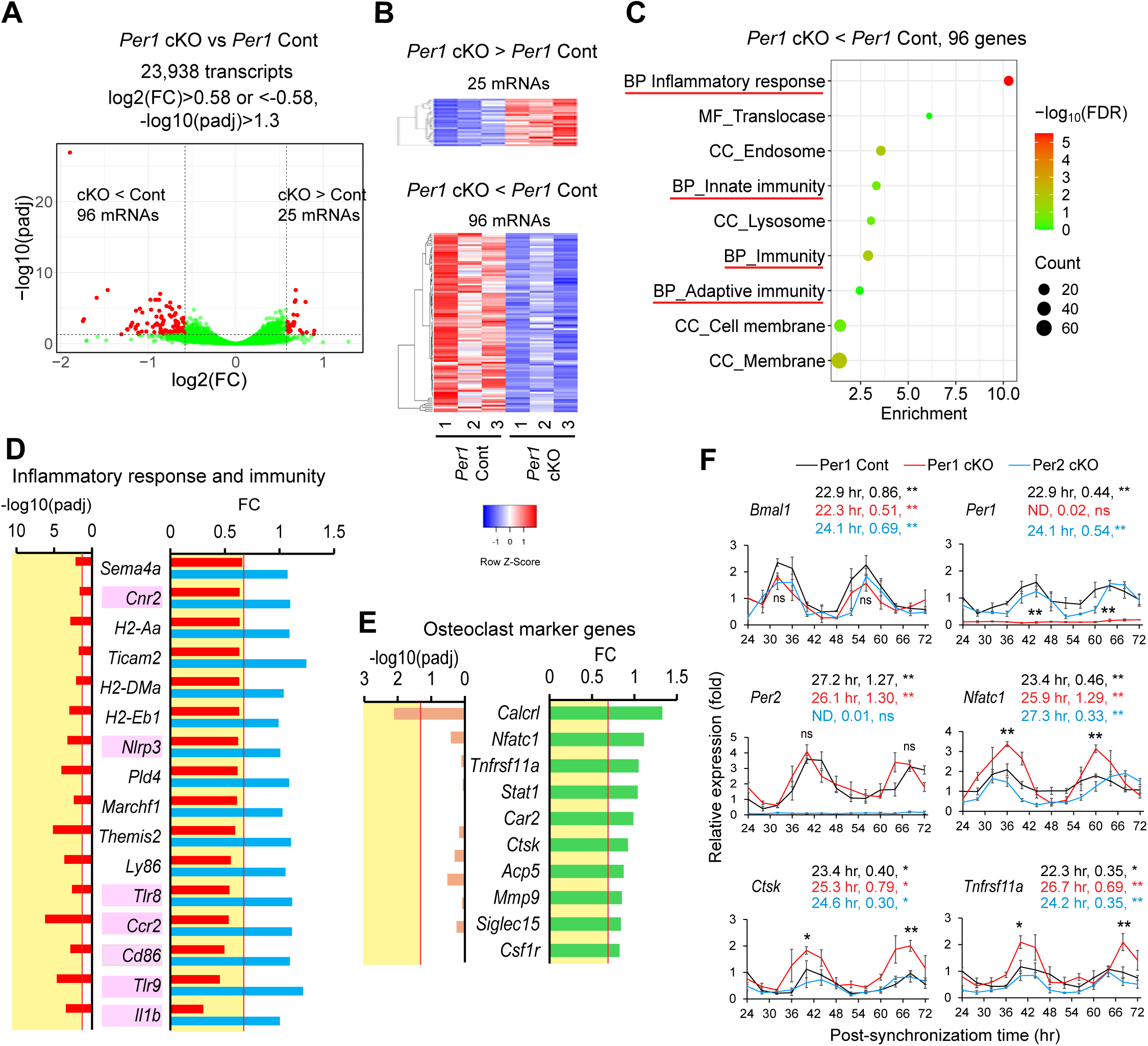
*Per1* cKO downregulated inflammatory genes in male osteoclasts. **A.** A volcano plot demonstrating differentially expressed genes between *Per1* cKO and Cont osteoclasts. FC and padj indicate fold change and adjusted p value, respectively. **B.** A heatmap displaying up- or downregulated genes in *Per1* cKO osteoclasts compared with *Per1* Cont osteoclasts. **C.** Gene ontology analysis of downregulated genes in *Per1* cKO osteoclasts compared with *Per1* Cont osteoclasts. Pathways related to inflammation and immunity are underlined in red. **D.** A list of the genes that belong to the four pathways underlined in red in (**C**). Red and blue bars indicate *Per1* cKO and *Per2* cKO osteoclasts, respectively. The areas of FC < 0.67 or padj > 0.05 (or –log10[padj] > 1.3) are highlighted in yellow in (**D**) and (**E**). **E.** A list of representative osteoclasts marker genes detected with RNA-seq. **F.** Relative expression levels of circadian regulators and osteoclast markers in synchronized osteoclasts comparing three genotypes. The value of *Per1* Cont cells at 24 hr was defined as 1.0 in each graph. The period (hr) and amplitude of each genotype are shown above each graph. ** p < 0.01 and ns for not significant with the Cosinor analysis of circadian rhythmicity above each graph. ND indicates not detected. ** p < 0.01, * p < 0.05, and ns with unpaired two-tailed t-test comparing peak levels of *Per1* cKO and Cont osteoclasts embedded in each graph. Male osteoclasts were used in all data. (**A**)–(**E**) are based on biological triplicates, whereas (**F**) is based on biological triplicates with technical triplicates each.

**Figure 4.**
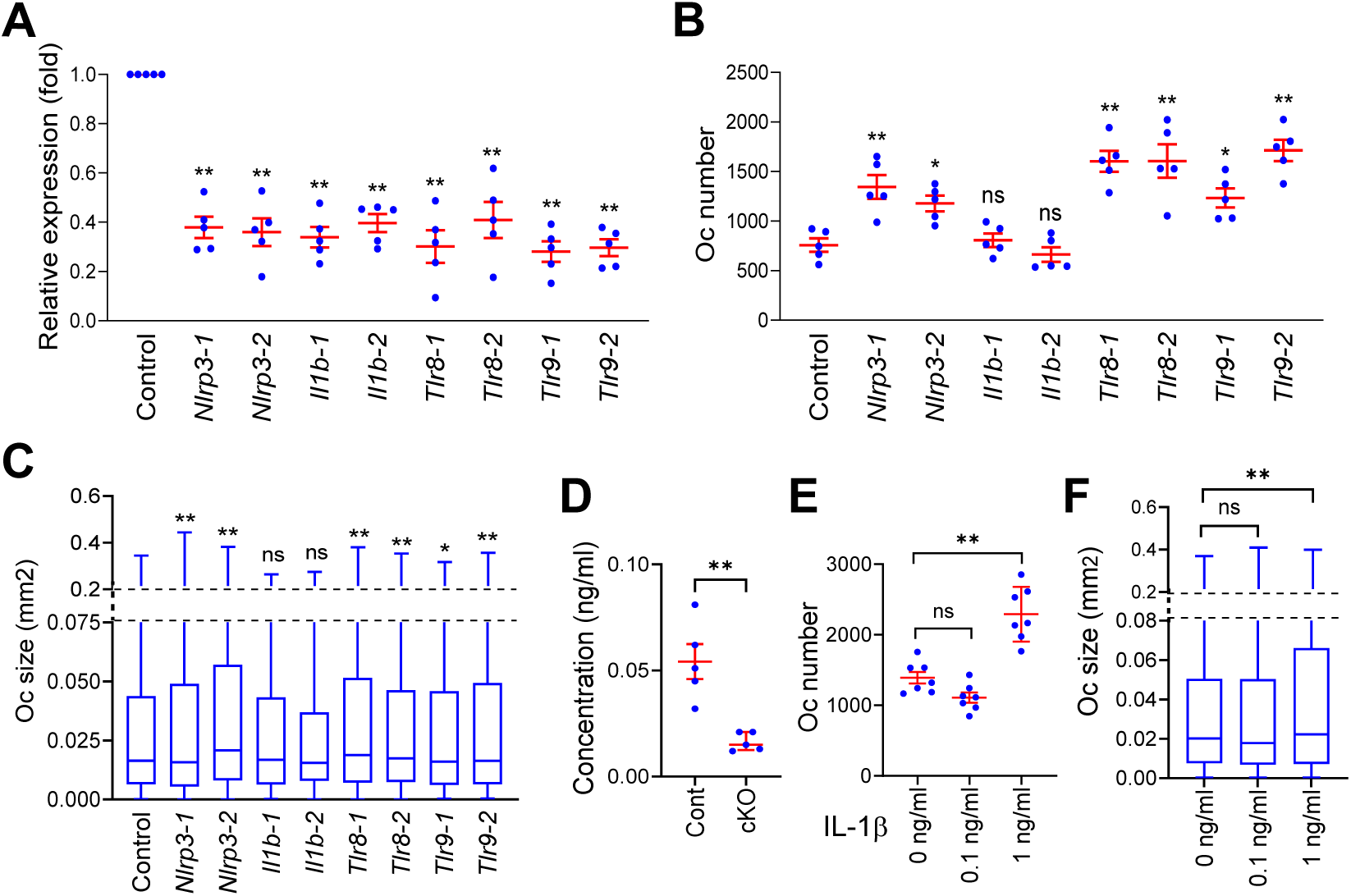
*Nlrp3*, *Tlr8*, and *Tlr9* inhibited osteoclastogenesis in vitro. **A.** Relative expression levels of the indicated genes after KD with two independent siRNA sequences each. The value with Control siRNA was defined as 1.0. n = 5 biological replicates with technical triplicates each. **B.** The numbers of osteoclasts in a well of a 48-well plate after KD of the indicated genes. n = 5. **C.** Size distributions of osteoclasts on day 6. n = 4,200 osteoclasts (600 osteoclasts/well x 7 wells of biologically independent experiments). **D.** Concentrations of IL-1β in the osteoclast supernatant quantified with ELISA. n = 5. **E.** The numbers of osteoclasts in a well of a 48-well plate after culture with IL-1β. n = 7. **F.** Size distributions of osteoclasts after culture with IL-1β. n = 4,200 (600 osteoclasts/per well x 7 wells of biologically independent experiments). All panels except for (**D**) used male *Per1* Cont osteoclasts. Mean ± SEM is shown. ** p < 0.01, * p < 0.05, and ns for not significant with unpaired two-tailed t-test in comparison to control samples in (**A**)–(**D**) and two-way ANOVA with Tukey’s method of multiple comparisons in (**E**) and (**F**).

The 16 downregulated genes included seven genes known to affect bone mass and osteoclastogenesis (**Figure 3D**, highlighted in pink, and **Table 1**). Although germline KO was reported for some genes (*Ccr2*, *Cd86*, and *Tlr9*), none of them have been studied using osteoclast-specific KO mice, making it difficult to compare their mouse phenotypes with those in the current study. In vitro studies of these genes showed that they were not simple inhibitors of osteoclastogenesis, as might be expected from the *Per1* cKO phenotypes. While *Ccr2* promotes osteoclastogenesis, *Il1b*, *Nlrp3*, and *Tlr9* have dual functions (inhibition and promotion) depending on the context. The decreased bone mass and increased osteoclastogenesis observed in our study may have resulted from cumulative effects of multiple dysregulated genes functioning at different stages of osteoclastogenesis.

**Table 1.** The roles of seven downregulated genes in osteoclastogenesis.

| **Gene** | **Relevance to osteoclastogenesis** | **Ref** |
| --- | --- | --- |
| *Ccr2* | C-C motif chemokine receptor 2. Increased bone mass and decreased osteoclasts in *Ccr2*<sup>-/-</sup> mice. Inhibited osteoclastogenesis by *Ccr2*<sup>-/-</sup> cells in vitro. | 43,44 |
| *Cd86* | Decreased bone mass and increased osteoclasts in *Cd80*<sup>-/-</sup>;*Cd86*<sup>-/-</sup> mice. Normal osteoclastogenesis by *Cd80*<sup>-/-</sup>;*Cd86*<sup>-/-</sup> monocytes in vitro. CD80 and CD86 are closely related. | 45 |
| *Cnr2*<br>(*CB2*) | Cannabinoid receptor 2. Uncertain roles. A study showed decreased bone mass and increased osteoclasts in *CB*<sup>-/-</sup> mice, while another indicated normal bone mass. | 46,47 |
| *Il1b* | IL-1β. Generally pro-osteoclastogenesis but can be anti-osteoclastogenesis depending on the culture conditions in vitro. | 27,28 |
| *Nlrp3* | NOD-, LRR- and pyrin domain-containing protein 3a. Inhibits osteoclastogenesis under physiological conditions in vitro but promotes it during infection. | 25,26 |
| *Tlr8* | Toll-like receptor 8. Binds to bacterial and viral single-stranded RNA. Promotes osteoclastogenesis when stimulated by a ligand in vitro. Stage-specific effect on osteoclastogenesis (below) is not known. | 48 |
| *Tlr9* | Recognizes DNA of bacteria and viruses and activates inflammation. Increased osteoclastogenesis and bone resorption in *Tlr9*<sup>-/-</sup> mice as a secondary effect of systemic inflammation. Inhibits or promotes osteoclastogenesis depending on the stage of differentiation. | 40,41,49 |

### Osteoclast marker genes were temporarily upregulated in circadian manners in *Per1* cKO cells

RNA-seq did not reveal expected upregulation of osteoclast marker genes in *Per1* cKO cells despite enhanced osteoclastogenesis (**Figure 3E**). We tested whether this was due to the cells’ circadian rhythms not being synchronized, which masked their temporary upregulation. To test this possibility, we synchronized the circadian rhythms by incubating the cells in 50% horse serum for 1 hr and then harvested them every 4 hr over a 48-hr period. The circadian expression patterns of *Bmal1*, *Per1*, and *Per2* in *Per1* Cont cells were verified using Cosinor analysis (**Figure 3F**, p < 0.01 for each gene, and **Supplementary Figure 5C**). In addition, the patterns of *Bmal1* and *Per2* expression were anti-phasic in both *Per1* cKO and Cont cells, validating successful synchronization. Moreover, *Per1* expression was substantially decreased in *Per1* cKO cells, as expected. Importantly, *Nfatc1*, *Ctsk*, and *Tnfrsf11a* exhibited circadian expression, with peak levels upregulated by *Per1* cKO, indicating that the transient upregulation of these genes in *Per1* cKO cells was obscured by the presence of cells at mixed circadian phases when the cells were not synchronized.

### *Per1* downregulated inflammasome pathway genes

*Nlrp3, Il1b*, *Tlr8*, and *Tlr9* are involved in the inflammasome signaling pathway, which promotes osteoclastogenesis.^24^ NLRP3, one of the main components of the inflammasome complex, activates caspase-1, leading to the cleavage of pro-IL-1β and pro-IL-18 to produce mature IL-1β and IL-18, increasing osteoclastogenesis.^24^ NLRP3 promotes osteoclastogenesis under inflammatory conditions created with LPS at 100 ng/ml; however, it inhibits osteoclastogenesis via pyroptosis under non-inflammatory conditions.^25,26^ In our case, LPS was undetectable (< 0.1 pg/ml) in the culture supernatants of *Per1* cKO and Cont cells on days 0 and 5, suggesting that NLRP3 functioned as an inhibitor of osteoclastogenesis. To verify this interpretation, we depleted *Nlrp3* with two independent siRNA sequences to < 50% of the control level obtained with scrambled siRNA (**Figure 4A**). The KD increased the number and size of osteoclasts, promoting osteoclastogenesis, supporting our interpretation (**Figure 4B** and **C**).

*Il1b* is generally pro-osteoclastogenic but it can be anti-osteoclastogenic when added into the culture prior to the addition of RANKL.^27,28^ Our KD of *Il1b* did not affect the number or size of osteoclasts (**Figure 4A–C**), possibly due to the low concentration of IL-1β in the supernatant of osteoclasts. Exogenous IL-1β promotes the osteoclastogenesis of BMMs at > 0.5 ng/ml in vitro.^29^ However, its concentration in the culture medium of *Per1* Cont osteoclasts was < 0.1 ng/ml, as measured by ELISA (**Figure 4D**); even 0.1 ng/ml was insufficient to promote osteoclastogenesis (**Figures 4E** and **F**). The low concentration was consistent with the report by Alam et al., which found no detectable IL-1β in the supernatant.^26^ Thus, IL-1β appeared to be irrelevant in our assay. TLR9 promotes or inhibits osteoclastogenesis depending on the culture conditions, as detailed in the Discussion. In our case, the KD of *Tlr8* and *Tlr9* increased both the size and number of osteoclasts (**Figure 4A–C**).

### *Per1*-dependent circadian expression of *Nlrp3*, *Tlr8*, and *Tlr9*

The circadian expressions of *Nlrp3*, *Tlr8*, and *Tlr9* were known,^30^ but whether they were dependent on *Per1* or *Per2* was unclear. qRT-PCR of synchronized cells indicated that *Per1* Cont and *Per2* cKO cells demonstrated circadian expressions of these genes (p < 0.05), while the rhythmicity was lost in *Per1* cKO cells (**Figure 5A** and **Supplementary Figure 6**). This study included two negative control genes that were related to inflammasomes but not differentially expressed between *Per1* cKO and Cont cells: *Tlr3* (log2[FC] = −0.3, and −log10[padj] = 0.17) and *Pycard* (log2[FC] = −0.06, and −log10[padj] = 0.05). These genes also exhibited circadian expression, which was not disrupted by *Per1* KO, underscoring the specific regulation of *Nlrp3*, *Tlr8*, and *Tlr9* by *Per1*. In a complementary experiment, we transduced *Clock*, *Bmal1*, *Per1*, and *Per2* in various combinations into the macrophage cell line RAW264.7 because plasmid transfection with osteoclasts was inefficient. The transfection of *Clock* and *Bmal1* did not increase the expression levels of *Nlrp3*, *Tlr8*, or *Tlr9* compared to that with the empty vector when the same total amount of plasmids was used (**Figure 5B**). However, these genes were upregulated when *Per1*, but not *Per2*, was included. The expression levels of the two control genes (*Tlr3* and *Pycard*) were not affected by the transfections. Collectively, these results supported our interpretation of the *Per1*-specific upregulation of *Nlrp3*, *Tlr8*, or *Tlr9*. Whether PER1 controls these genes directly or indirectly through other genes regulated by circadian rhythms remains unknown.

**Figure 5.**
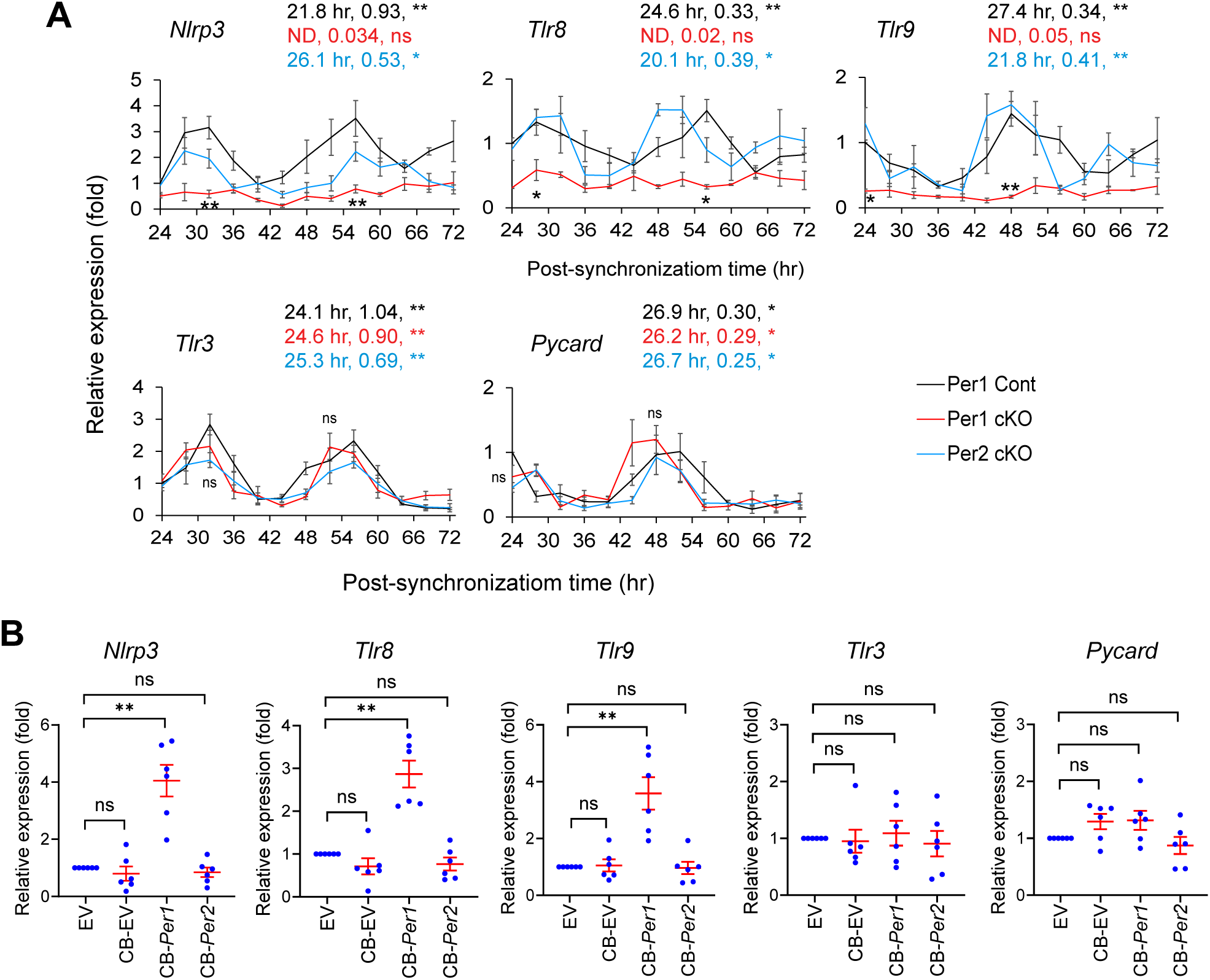
*Per1* controlled circadian expression of *Nlrp3*, *Tlr8*, and *Tlr9* in osteoclasts. **A.** Relative expression levels of the indicated genes in synchronized osteoclasts comparing *Per1* cKO, *Per2* cKO, and *Per1* Cont osteoclasts. The value of *Per1* Cont cells at 24 hr was defined as 1.0 in each graph. The period (hr) and amplitude of each genotype are shown above each graph. ** p < 0.01, * p < 0.05, and ns for not significant with the Cosinor analysis of circadian rhythmicity above each graph. ND indicates not detected. ** p < 0.01, * p < 0.05, and ns with unpaired two-tailed t-test comparing peak levels of *Per1* cKO and Cont osteoclasts embedded in each graph. **B.** Relative expression levels of the indicated genes after overexpression of circadian regulators in RAW264.7 cells. The value with EV was defined as 1.0. Abbreviations are as follows. EV: empty vector and CB: *Clock* and *Bmal1*. ** p < 0.01, * p < 0.05, and ns for not significant with two-way ANOVA with Tukey’s method of multiple comparisons. All experiments used male osteoclasts. Mean ± SEM is shown.

## Discussion

This study demonstrates that *Per1* cell-autonomously inhibits osteoclastogenesis in vitro, likely achieved by the upregulation of a group of inflammatory genes known to control osteoclastogenesis in context-dependent manners. These roles could explain the decreased bone mass in *Per1* cKO mice, suggesting the presence of the *Per1*–inflammatory genes–osteoclast axis. This axis appears to be influenced by sex because bone mass was not decreased in female *Per1* cKO mice. The underlying mechanism for this observation remains unknown although the roles of sex steroids in osteoclastogenesis have been well documented.^31^ Since the synthesis and metabolism of sex steroids mainly take place outside the tissues expressing *Cx3cr1*, the focus of the future research should be the crosstalk between the downstream of sex steroids and inflammatory genes in osteoclasts. In contrast to *Per1*, *Per2* cKO did not affect bone mass or osteoclastogenesis despite sharing a high similarity at the amino acid level (73.4%) with *Per1.* Our finding is the second example of *Per1*-specific regulation of inflammatory genes, following the example that we reported in macrophages, although the target inflammatory genes were different.^13^ The involvement of inflammatory genes was not discussed in the studies on *Bmal1* KO mentioned above.

We selected the *Cx3cr1* promoter to drive the *Cre* gene, instead of the *Ctsk* promoter because *Ctsk* is also expressed in periosteal mesenchymal stem cells, which can differentiate into osteoblasts,^32^ thereby compounding the interpretation of our results. However, the *Cx3cr1* is also expressed in the cells other than the monocyte/macrophage lineages, such as dendritic cells, natural killer (NK) cells, and some T cell subtypes.^33^ Depletion of *Per1* in these cells is also known to affect bone remodeling via secreted cytokines. For example, *Per1* KO in NK cells alters the peak level and timing of the circadian expression of interferon γ (IFN-γ) and cytolytic factors (perforin and granzyme B).^34^ *Per1* KD in helper T cells increases the synthesis of IFN-γ, IL-2, and TNF-α.^35^

Although these studies did not describe any bone phenotypes, it is possible that the cells indirectly affected bones because these cytokines are involved in bone remodeling and osteoporosis.^36–38^ Recent advances in single-cell spatial transcriptomics may enable us to analyze the contributions of each cell type. This approach would also allow us to verify whether the downregulated genes in the RNA-seq were reproducible in vivo, which is particularly significant for the dual-function genes *Nlpr3*, *Tlr9*, and potentially *Tlr8*.

TLR8 and TLR9 belong to the pattern recognition receptors in the innate immune system, which are localized on the membranes of endosomes, lysosomes, and endolysosomes.^39^ TLR8 binds to single-stranded RNA, including viral RNA, whereas TLR9 binds to unmethylated CpG-containing DNA, which is more abundant in viral and bacterial DNA than in mammalian DNA. Binding these molecules leads to the production of pro-inflammatory cytokines (such as IL-1β and TNF-α) and type I interferons (such as IFN-α and IFN-β), which are generally pro-osteoclastogenic. TLR9 inhibits osteoclastogenesis when it is activated in the presence of M-CSF and RANKL at the start of culture, but promotes osteoclastogenesis when activated after the cells have been primed by M-CSF and RANKL for three days.^40,41^ The inhibition of osteoclastogenesis by TLR9 has been explained by the enhanced degradation of the c-fos protein, an essential inducer of osteoclastogenesis.^42^ However, the relevance of this culture condition-dependent difference in osteoclastogenesis to physiological osteoclastogenesis in vivo remains unclear. In the present study, TLR9 was not explicitly activated, although the possibility of unintentional activation could not be excluded. RANKL was added to the culture medium from day 2, and *Tlr9* siRNA was introduced on day 4, leading to enhanced osteoclastogenesis. This promotion likely reflects the role of TLR9 at the baseline (non-activated) level, which could change once it is activated under an inflammatory condition. Although TLR8 has not been characterized in detail, it likely has similar roles due to the shared downstream signaling pathways.

The involvement of several genes relevant to inflammasomes in osteoclastogenesis raises the possibility of the inflammatory reactions of osteoclasts as a part of the physiological *Per1*-mediated circadian regulation of bone remodeling. This precondition would be amplified when inflammasomes are activated by microorganisms or cell debris. This may impact inflammatory bone diseases influenced by circadian rhythms, such as rheumatoid arthritis and osteoarthritis, where osteoclast-mediated bone destruction is a significant factor.^6^ Further mechanistic studies on *Per1*-specific circadian regulation of inflammatory genes could lead to the identification of novel therapeutic options for the inflammatory bone diseases, in addition to non-inflammatory predispositions to osteoporosis caused by circadian disruption, as broader implications of this research.

## Supporting information

Graphical abstract

## Acknowledgments

We thank Drs. Bryce Binstadt and Michelle L. Gumz for *Cx3cr1*-*Cre* mice and *Per1^fl/fl^* mice, respectively. We are grateful to the Comparative Pathology Shared Resource and the Minnesota Dental Research Center for Biomaterials and Biomechanics.

## Author contributions

Nobuko Katoku-Kikyo (investigation, writing - review and editing), Elizabeth K. Vu (investigation, writing - review and editing), Samuel Mitchell (investigation, writing - review and editing), Ismael Y. Karkache (investigation, writing - review and editing), Elizabeth W. Bradley (conceptualization, funding acquisition, investigation, writing - review and editing), and Nobuaki Kikyo (conceptualization, funding acquisition, investigation, writing - original draft, writing - review and editing).

## Data Availability

RNA-seq data have been deposited to Gene Expression Omnibus (GEO) under the accession number of GSE283894 (*Per1* cKO and control) and GSE292534 (*Per2* cKO and control).

## Funding Statement

E.W.B. was supported by the NIH (R21AR084530). N.K was supported by the NIH (R01GM137603) and Regenerative Medicine Minnesota (RMM 072523 DS 002). The content is solely the responsibility of the authors and does not necessarily represent the official views of the NIH.

## Conflict of interest

The authors declare no conflict of interest.

## Notes

### Competing Interest Statement

The authors have declared no competing interest.

### Summary of Updates

Text and figures were revised in response to journal reviewers' comments.

