## Supplementary Materials for "The circadian regulator PER1 inhibits osteoclastogenesis by activating inflammatory genes"

**List of Supplementary Materials in PDF**

Supplementary Figure 1–6

Supplementary Figure Legends

Supplementary Table 1–3

Supplementary Fig 1

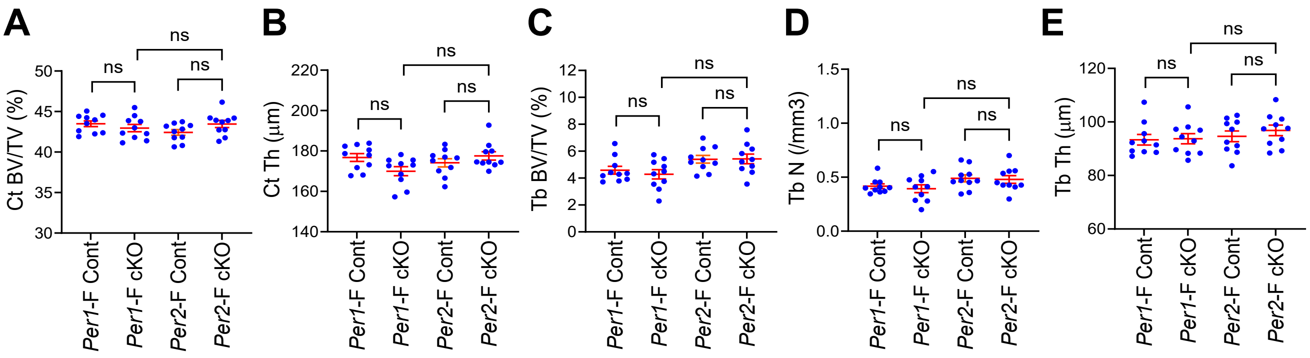

Supplementary Fig 2

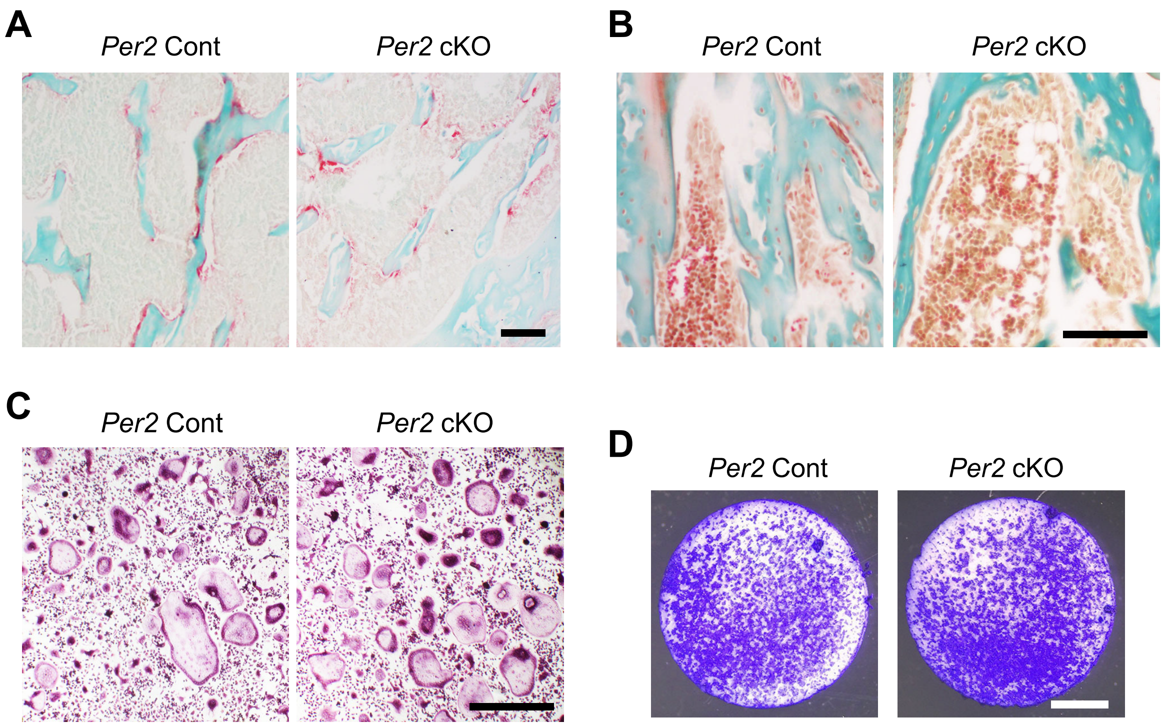

Supplementary Fig 3

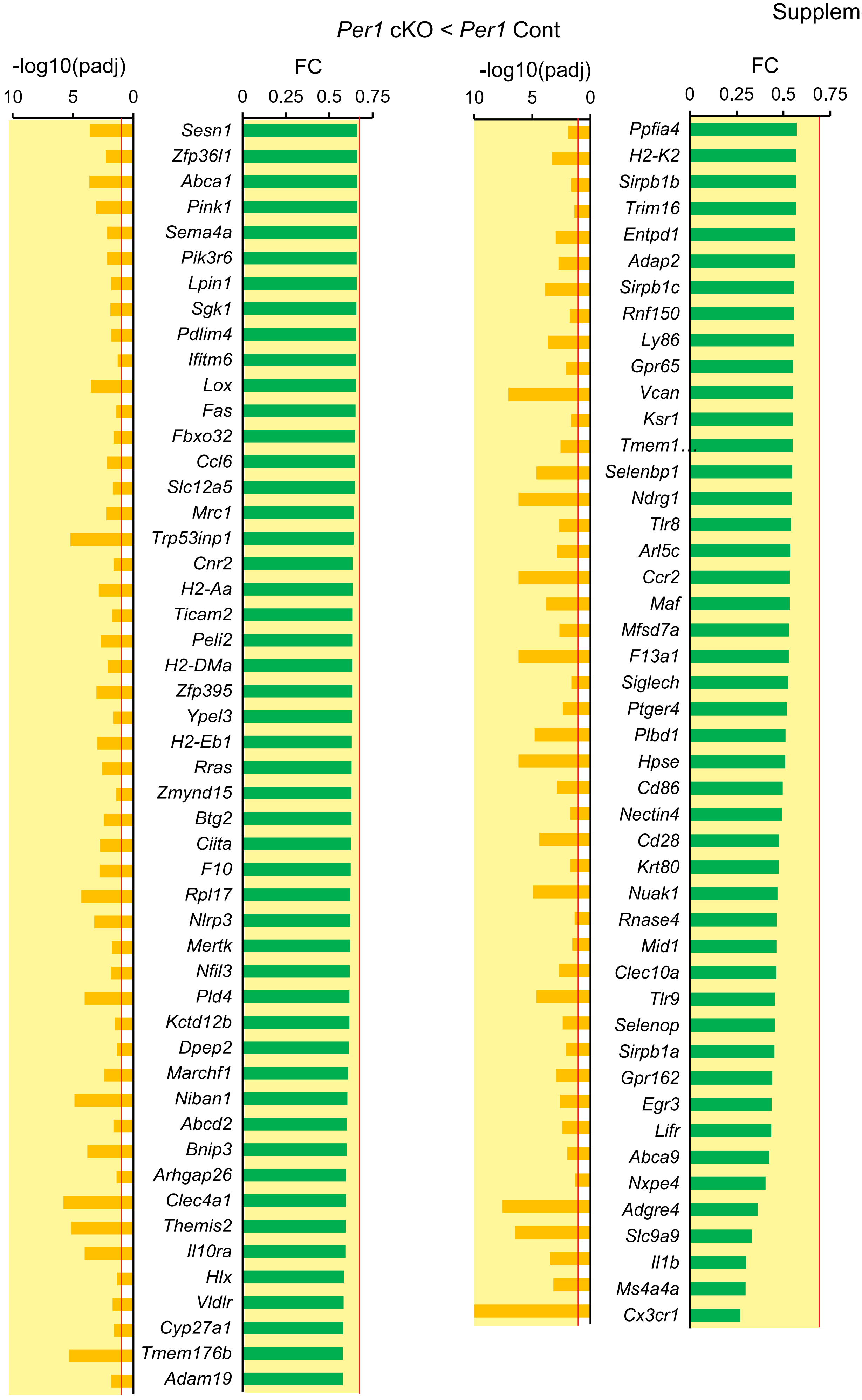

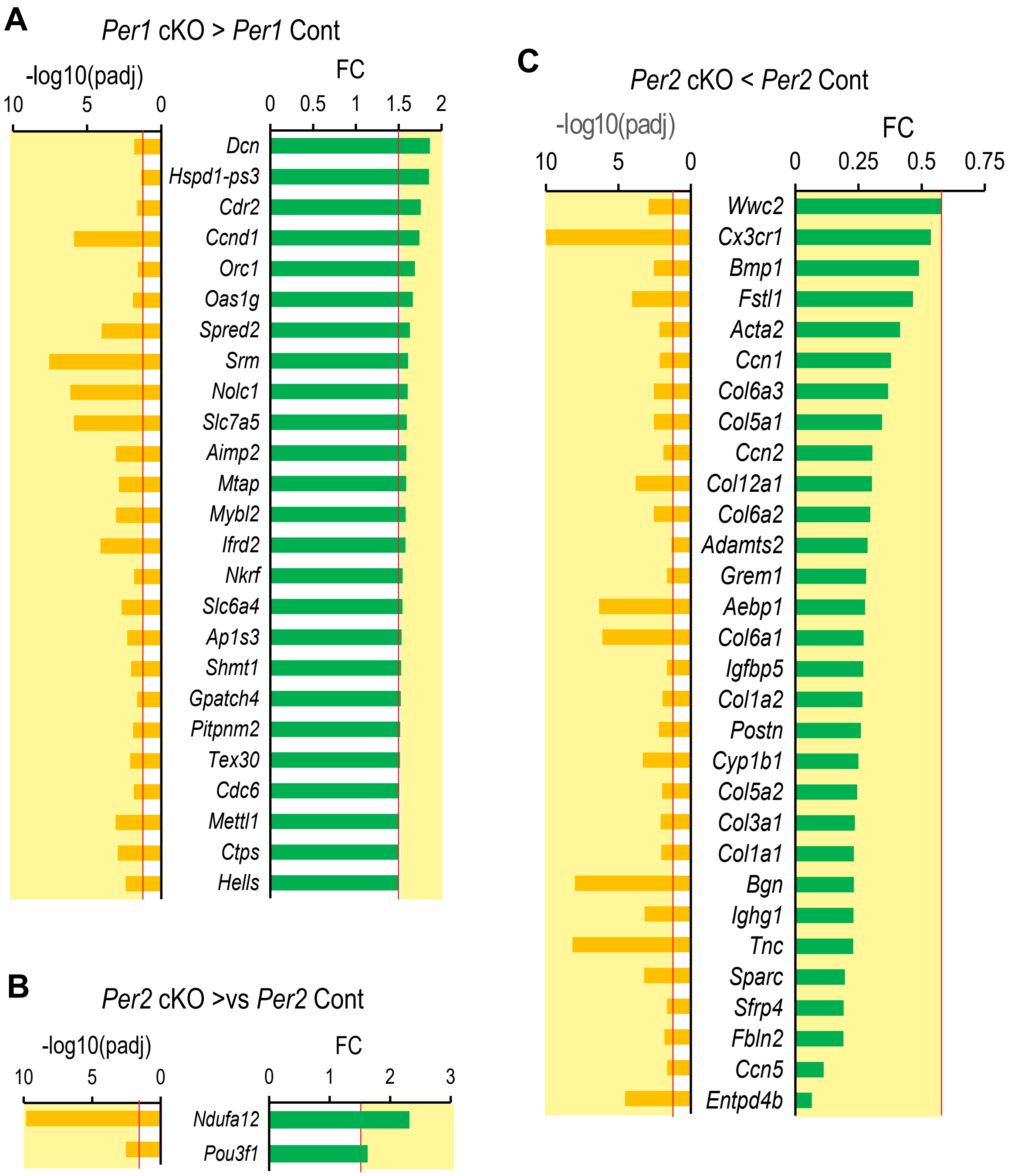
A
Per1 cKO > Per1 Cont
-log10(padj)
10 5 0
FC
0 0.5 1 1.5 2
Dcn
Hspd1-ps3
Cdr2
Ccnd1
Orc1
Oas1g
Spred2
Srm
Nolc1
Slc7a5
Aimp2
Mtap
Mybl2
Ifrd2
Nkrf
Slc6a4
Ap1s3
Shmt1
Gpatch4
Pitpnm2
Tex30
Cdc6
Mettl1
Ctps
Hells
B
Per2 cKO >vs Per2 Cont
-log10(padj)
10 5 0
FC
0 1 2 3
Ndufa12
Pou3f1
C
Per2 cKO < Per2 Cont
-log10(padj)
10 5 0
FC
0 0.25 0.5 0.75
Wwc2
Cx3cr1
Bmp1
Fstl1
Acta2
Ccn1
Col6a3
Col5a1
Ccn2
Col12a1
Col6a2
Adamts2
Grem1
Aebp1
Col6a1
Igfbp5
Col1a2
Postn
Cyp1b1
Col5a2
Col3a1
Col1a1
Bgn
Ighg1
Tnc
Sparc
Sfrp4
Fbln2
Ccn5
Entpd4b

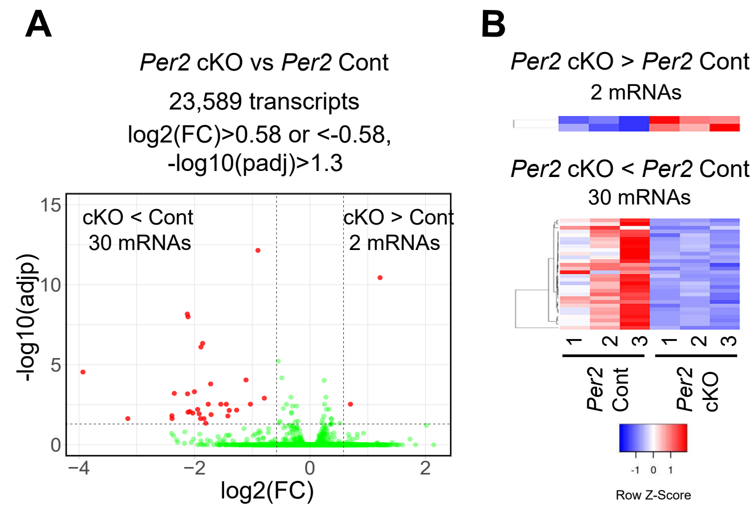
A
*Per2* cKO vs *Per2* Cont
23,589 transcripts
log2(FC)>0.58 or <-0.58,
-log10(padj)>1.3
cKO < Cont
30 mRNAs
cKO > Cont
2 mRNAs
-log10(adjp)
log2(FC)
B
*Per2* cKO > *Per2* Cont
2 mRNAs
*Per2* cKO < *Per2* Cont
30 mRNAs
*Per2* Cont
*Per2* cKO
Row Z-Score

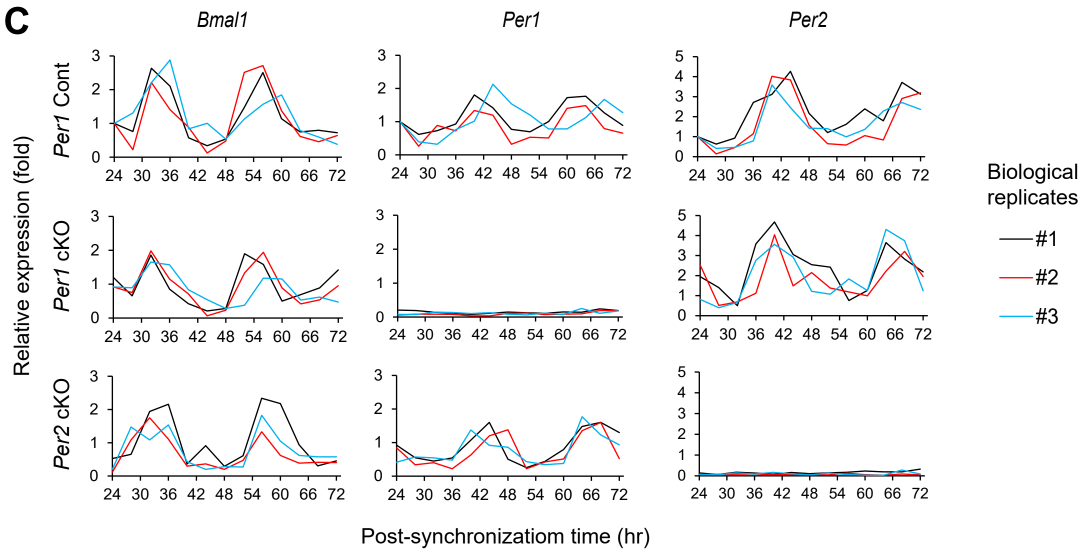
C
*Bmal1*
*Per1*
*Per2*
*Per1* Cont
*Per1* cKO
*Per2* cKO
Relative expression (fold)
Post-synchronizatiom time (hr)
Biological replicates
#1
#2
#3

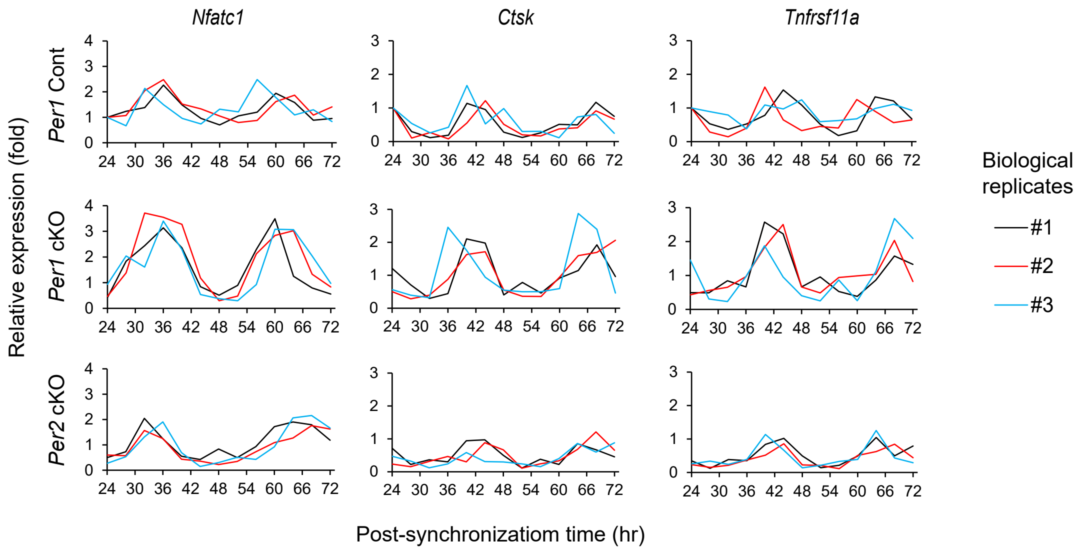
*Nfatc1*
*Ctsk*
*Tnfrsf11a*
*Per1* Cont
*Per1* cKO
*Per2* cKO
Relative expression (fold)
Post-synchronizatiom time (hr)
Biological replicates
#1
#2
#3

Supplementary Fig 6

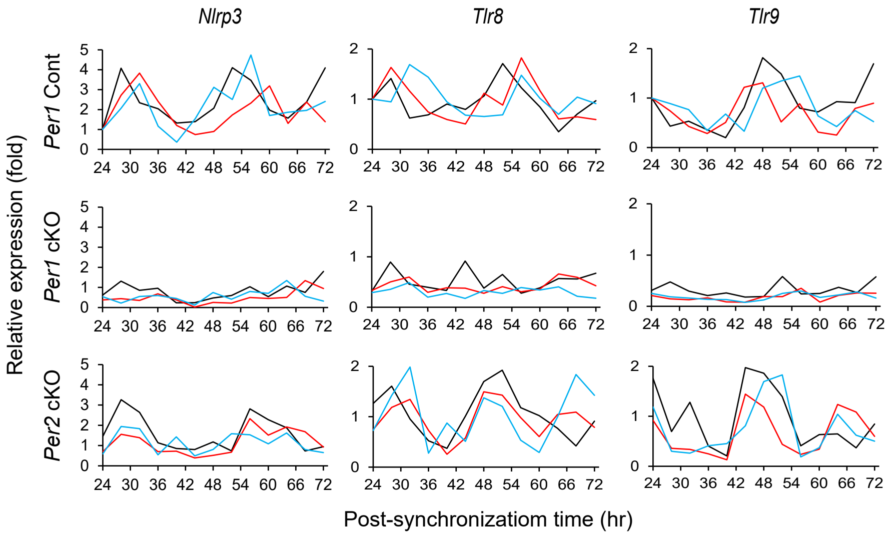


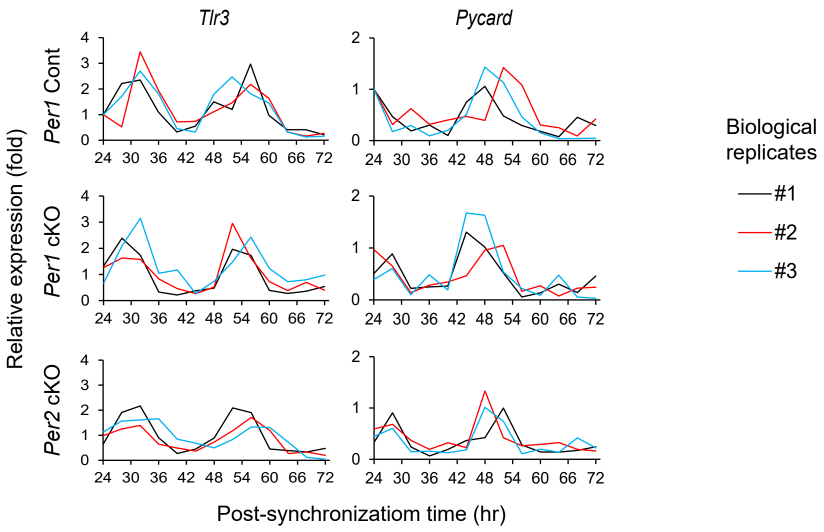

**Supplementary figure legends**

**Supplementary Figure 1. Bone mass of the female femurs was not affected by *Per1* or *Per2* cKO.**

**A**–**E**. Quantification of the cortical bone volume/total volume ratio (**A**), cortical thickness (**B**), trabecular bone volume/ total volume ratio (**C**), trabecular number (**D**), and trabecular thickness (**E**) comparing 12-week-old female mice. n=10.

ns for not significant with two-way ANOVA with Tukey's method of multiple comparisons.

**Supplementary Figure 2. *Per2* cKO in male mice did not affect osteoclastogenesis or osteoblastogenesis.**

**A.** TRAP staining of the proximal tibial sections comparing *Per2* cKO and Cont mice. Bar, 100 μm.

**B.** Masson's trichrome staining of the proximal tibial sections. Bar, 100 μm.

**C.** TRAP staining of osteoclasts on day 6. Bar, 500 μm.

**D.** Bone resorption assay stained with Toluidine blue. Bar, 2 mm.

Histological sections and osteoclasts were prepared from 12-week-old male mice.

**Supplementary Figure 3. A list of genes downregulated by *Per1* cKO in osteoclasts.**

All data are based on n = 3 of male cells. The areas of FC < 0.67 or padj > 0.05 (or –log10[padj] > 1.3) are highlighted in yellow.

**Supplementary Figure 4. Lists of genes dysregulated by cKO of *Per1* or *Per2* in osteoclasts.**

**A.** Genes upregulated by *Per1* cKO.

**B.** Genes upregulated by *Per2* cKO.

**C.** Genes downregulated by *Per2* cKO.

All data are based on n = 3 of male cells. The areas of FC < 0.67, FC > 1.5, or padj > 0.05 (or –log10[padj] > 1.3) are highlighted in yellow.

**Supplementary Figure 5. Analysis of dysregulated genes by *Per1* or *Per2* cKO in osteoclasts.**

**A.** A volcano plot demonstrating differentially expressed genes between *Per2* cKO and Cont osteoclasts. n = 3 male osteoclasts in (**A**) and (**B**).

**B.** A heatmap displaying up- or downregulated genes in *Per2* cKO osteoclasts compared with *Per2* Cont osteoclasts.

**C.** Breakdown of the data shown in Figure 3F into individual biological replicates. Each graph represents the average of technical triplicates of qPCR.

**Supplementary Figure 6. Circadian expression of inflammatory genes in osteoclasts.**

Breakdown of the data shown in Figure 5A into individual biological replicates. Each graph represents the average of technical triplicates of qPCR.

**Supplementary Table 1. Sequences of genotyping primers and the sizes of the PCR products**

| Gene | Sequence |
|---|---|
| *Cx3cr1* forward for the wild type allele | CCTCAGTGTGACGGAGACAG |
| *Cx3cr1* forward for the *Cre* allele | GACATTTGCCTTGCTGGAC |
| *Cx3cr1* reverse | GCAGGGAAATCTGATGCAAG |
| Sizes of PCR products of *Cx3cr1* | 302 bp for the wild type allele and 380 bp for the *Cre* allele |
| *Per1* forward | ATGAAGGTGGATAGGCTAGGGC |
| *Per1* reverse | GCCTTACCTTTCATCTACATCCTGG |
| Sizes of PCR products of *Per1* | 459 bp for the wild type allele and 539 bp for the floxed allele |
| *Per2* forward | GGGACCTGACCCATCATTCT |
| *Per2* reverse for the wild type allele | TAGGCTTCACCACAGGGTTC |
| *Per2* reverse for the floxed allele | GAACTTCGGAATAGGAACTTCG |
| Sizes of PCR products of *Per2* | 248 bp for the wild type allele and 147 bp for the floxed allele |

**Supplementary Table 2. Sequences of qPCR primers**

| Gene | Forward | Reverse |
|---|---|---|
| *Gapdh* | TGCACCACCAACTGCTTAG | GATGCAGGGATGATGTTC |
| *Bmal1* | CAACCCATACACAGAAGCAAAC | CATCTGCTGCCCTGAGAATTA |
| *Per1* | CCTGGAGGAATTGGAGCATATC | CCTGCCTGCTCCGAAATATAG |
| *Per2* | CAAAGCTGACGCACACAAAG | TTAGCCTTCACCTGCTTCAC |
| *Il1b* | CCACCTCAATGGACAGAATATCA | CCCAAGGCCACAGGTATTT |
| *Nlrp3* | GTTCTGAGCTCCAACCATTCT | CACTGTGGGTCCTTCATCTTT |
| *Tlr8* | GTAACGCACCGTCTAGGATTT | TTCAGCTCACTTTCCTCTGTG |
| *Tlr9* | TGGACGGGAACTGCTACTA | CAGAGACAGATGGGTGAGATTG |
| *Tlr3* | ACCTCCAGAAGAACCTCATAAC | GAACGGATTGAAGCGCATATC |
| *Pycard* | ACCAGCCAAGACAAGATGAG | CCATCACCAAGTAGGGATGTATT |

### Supplementary Table 3. Target DNA sequences of siRNAs

| Gene | Target DNA sequence |
|---|---|
| *Non-targeting control* | TGGTTTACATGTCGACTAA |
| *Il1b-1* | GAGGACATGAGCACCTTCTTT |
| *Il1b-2* | GCAGGCAGTATCACTCATTGT |
| *Nlrp3-1* | GCCATGTGGAGATCCTAGGTT |
| *Nlrp3-2* | GGAGTTCTTCGCTGCTATGTA |
| *Tlr8-1* | GAACCAGTGTTACAGTACTCA |
| *Tlr8-2* | GGATTTAACCAACAACAGACT |
| *Tlr9-1* | GGACAGGTGTAAGAACTTCAA |
| *Tlr9-2* | GTTCAGTGAGCTACCACAGTT |
