## Supplementary material for "The circadian regulator PER1 inhibits osteoclastogenesis by activating inflammatory genes": Graphical abstract

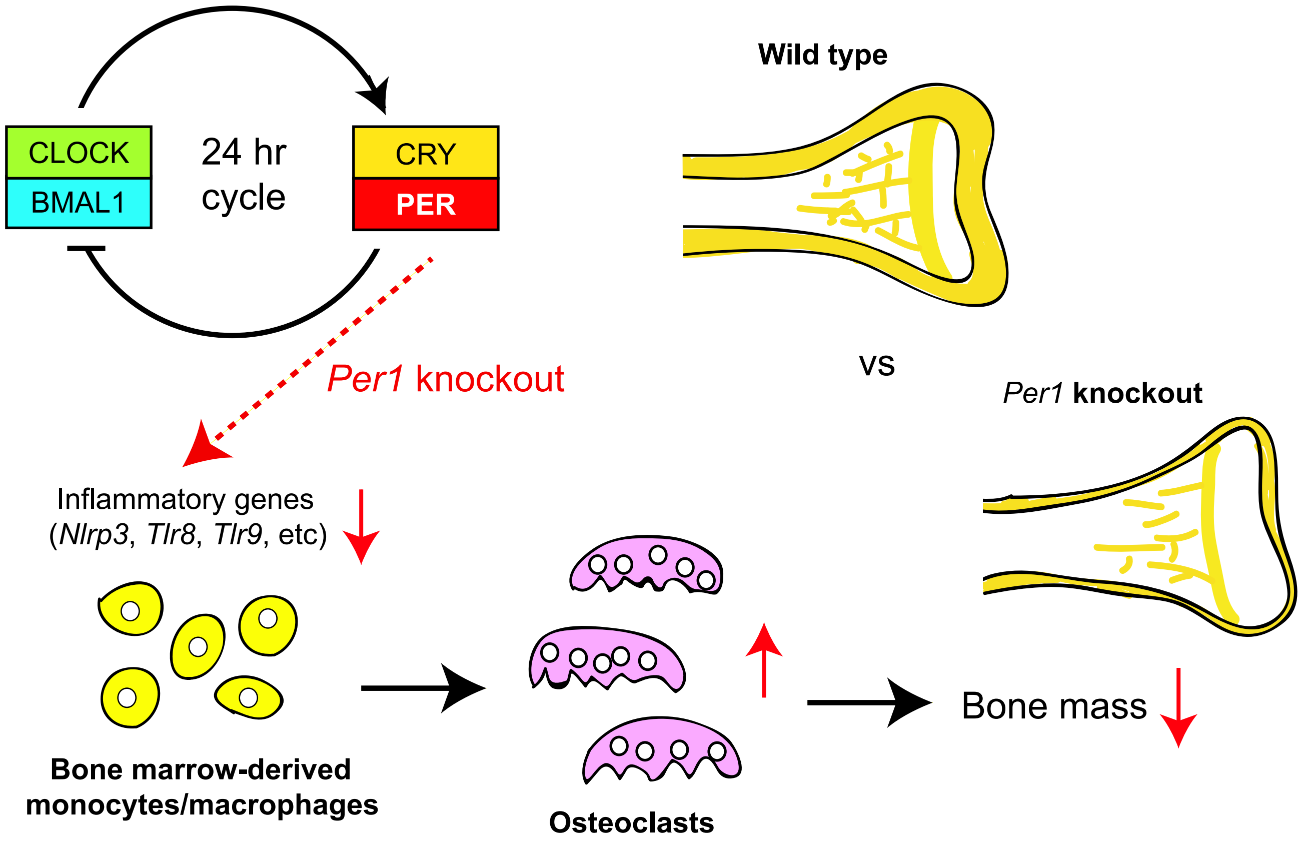

**Graphical abstract**

The mammalian CLOCK/BMAL1 heterodimer activates hundreds of target genes including *Cry* and *Per*, whose proteins inhibit CLOCK/BMAL1, thus generating circadian rhythms with a 24-hr period. When *Per1* is knocked out, inflammatory genes are downregulated in bone marrow-derived monocytes and macrophages, which promotes osteoclastogenesis and decreases bone mass.
